# Nanoscale patterning of *in vitro* neuronal circuits

**DOI:** 10.1101/2021.12.16.472887

**Authors:** José C. Mateus, Sean Weaver, Dirk van Swaay, Aline F. Renz, Julian Hengsteler, Paulo Aguiar, János Vörös

## Abstract

Methods for patterning neurons *in vitro* have gradually improved and are used to investigate questions difficult to address *in* or *ex vivo*. Though these techniques guide axons between groups of neurons, multiscale control of neuronal connectivity, from circuits to synapses, is yet to be achieved *in vitro*. As studying neuronal circuits with synaptic resolution *in vivo* poses significant challenges, an *in vitro* alternative could serve as a testbed for *in vivo* experiments or as a platform for validating biophysical and computational models. In this work we use a combination of electron beam and photolithography to create polydimethylsiloxane (PDMS) structures with features ranging from 150 nanometers to a few millimeters. Leveraging the difference between average axon and dendritic spine diameters, we restrict axon growth while allowing spines to pass through nanochannels to guide synapse formation between small groups of neurons (i.e. nodes). We show this technique can be used to generate large numbers of isolated feed-forward circuits where connections between nodes are restricted to regions connected by nanochannels. Using a genetically encoded calcium indicator in combination with fluorescently tagged post synaptic protein, PSD-95, we demonstrate functional synapses can form in this region. Although more work needs to be done to control connectivity *in vitro*, we believe this is a significant step in that direction.

---

The brain controls the body via billions of neurons dedicated to information processing. In turn, each neuron can establish thousands of synapses, expanding the complexity of neuronal circuits by orders of magnitude. Synapses connect neurons in a structural network that can be termed as the synaptome^1^, analogous to the connectome that defines the functional network. Notably, synapses are remarkably plastic structures with modifications in their number, size, or strength being associated with higher-order functions, such as learning or memory^2,3^. Understandably, there is a great interest in developing tools and methods for monitoring synapse formation, maintenance, or plasticity in long-term experiments. Ideally, these could allow for a continuous synaptome and connectome mapping of neuronal circuits, with a wide range of applications in studies of neuronal physiology and pathology.

Numerous limitations hamper the ability to experimentally probe neurons, and synapses, *in vivo*; however, *in vitro* models can serve as unique experimental tools to unveil universal mechanisms of neuronal circuits. Historically, studies using low-density neuronal cultures have revealed fundamental properties of synaptic plasticity, such as spike timing-dependent plasticity (STDP)^4,5^. However, the morphological and functional complexity of these seemingly random neuronal networks makes the systematic study of their connectivity challenging. So-called “brain-on-a-chip” technologies have improved in terms of high-throughput capabilities in the last few years, particularly for disease modeling and pharmacological testing (reviewed^6,7,8^). For the interrogation of neuronal circuits, these devices may also be engineered to gain better control over neuronal network topology and connectivity^9^. This allows for the study of small-scale connectomes, which is nearly impossible to achieve *in vivo* for vertebrate models and conventional *in vitro* models^10^. Importantly, several studies link findings in engineered (“modular” or “node-like”) cultures^11,12,13^ to properties found at a larger scale and *in silico*^10,14^ validating these tools for the study of fundamental mechanisms of neuronal circuits. Despite their potential, until now, these tools have not been adapted to the study of small-scale synaptomes. With many disease models and pharmacological testing focusing on synaptic changes (for reviews see^15,16^), the ability to compartmentalize and interrogate these substructures is of fundamental importance.

Engineering neuronal network topology requires a combination of methods for controlled cell placement and growth. Several techniques for controlling these parameters *in vitro* have been proposed in the literature (reviewed in^17,18^). The most widely used methods create patterns of cell-adhesive and/or cell-repellent promoting zones on 2D substrates (chemical patterning; e.g., microcontact printing)^13,19,20,21,22,23^ or create 3D structures that physically confine and guide neurons (physical patterning; e.g., microfluidics).^9,24,25,26^ Typically, both approaches take advantage of conventional soft-lithography for stamping/structuring the desired patterns at the microscale (e.g., microspots/microwells) by using biocompatible silicones^27^, most often poly(dimethylsiloxane) (PDMS),^28^ which are replicated from a photolithographically patterned master mold. In particular, PDMS devices that create microfluidic environments to compartmentalize neurons and guide neurites through subcellular microchannels, are widely used across a range of neuroscience fields and as the main tool for the structuring of a brain-on-a-chip (reviewed in^7,29^). Critically, these methods have allowed researchers to build mesoscale circuits with varying levels of control over circuit topology, but neurons’ positioning and connections remain highly random.

The dimensions of critical morphological traits for the establishment of synapses range from tens of nanometers (e.g., dendritic spines) to millimeters (axonal length). To faithfully control microcircuit formation, advances for *in vitro* models need to encompass all these patterning scales. However, due to the inherent fabrication and physiological difficulties, nanoscale guidance has, to our knowledge, so far been neglected. State-of-the-art studies have used microscale features to control neuron position,^20,30^ axon guidance,^7,24^ or both^9,23,31,32,33,34,35^ but without control over synapse formation. Consequently, there are no dedicated tools/methods to precisely impose where, and between which neurons, synapses form - a key limitation in the study of neuronal microcircuits. New *in vitro* tools could allow for the control and probing of spine growth/synapse formation, thus facilitating the study of dependent mechanisms, such as synaptic scaling or plasticity.^3,11^ Ideally, these tools and methods should control neuron position and number, dendritic and axonal outgrowth, as well as synapse formation.

In general, progress in neuronal patterning has been substantial, but the current state-of-the-art lacks control over the precise positioning and connectivity of neurons needed for reproducible microcircuit formation and the study of synapses. Undoubtedly, there is a need to engineer advanced *in vitro* models at the nano-microscale that can recapitulate synapse formation in a well-controlled setting.^36,37^ Here, we detail new fabrication and cell culturing methods for advancing the neuroengineering field in these two shortcomings. Using nano-microscale patterning we present a new paradigm for the study of input-to-output (i.e., feed-forward) isolated microcircuits *in vitro*. Via fluorescent labeling of spine growth, synapse formation, and synaptic activity, we demonstrate design feasibility and an unprecedented degree of control over circuit connectivity. Moreover, these new *in vitro* models allow for the interrogation of neuronal microcircuits with several state-of-the-art tools, such as microelectrode arrays (MEAs) or genetically encoded sensors. Ultimately, the demonstrated ability to create isolated and precisely connected neuronal microcircuits, that can be easily probed and manipulated in a high-throughput manner, may have a great impact on bottom-up neuroscience.

## RESULTS and DISCUSSION

### Novel design for multiscale neuronal patterning

Most neuronal patterning methods use conventional photolithography as the feature limiting step of the fabrication and as such are restricted to minimum feature sizes of ∼2µm in at least two physical dimensions.^38^ While this is sufficient for isolating cell bodies (∼15µm), dendrites (∼2µm), and groups of axons (>0.5µm), (**Fig. 1A i-iii)** it is not possible to pattern individual axons, dendritic spines, or synaptic boutons (**Fig. 1A iii-v**). To recapitulate the organization of *in vivo* neuronal circuits, which demonstrate a high degree of structure at the synaptic level, techniques for neuron placement and axon guidance need to be combined with submicron features guiding synapse formation (**Fig. 1B i**). For excitatory synapses across many brain regions, the postsynaptic (i.e., receiving) element is a bulbous spine projected from the main dendrite body by a thin, ∼100nm diameter, spine neck which then comes in proximity, ∼20nm, to a presynaptic (i.e., sending) *en passant* bouton on the axon to form a synapse. While proximity alone is not enough to establish a synapse between an axon and a dendrite,^39^ it is a necessary condition. Though it is possible for segments of the axon to make thin protrusions (i.e., filopodia) with diameters thinner than 150nm, main axonal sections have a larger minimum diameter governed by the membrane periodic skeleton (e.g., actin rings). The axon diameter of rat hippocampal neurons *in vitro* is on average 450nm.^40,41^ By creating what can be considered submicron channels, which we will call nanochannels throughout the paper, with cross-sections well below the axon diameter but similar in size or larger than dendritic spine necks, we wish to impose where and between which groups of neurons synapses can form (**Fig. 1B ii, iii**). To validate this technique, we designed a neuronal circuit composed of three nodes (i.e., groups) of cells: one presynaptic and two postsynaptic. We decided on this circuit as it is one of the simplest to test while allowing for interesting features such as directional propagation of activity (i.e., feed-forward), multiple postsynaptic partners, and timing variation in signal propagation in addition to being a testbed platform for different types of plasticity (e.g., Hebbian-type or homeostatic). Throughout this work, we have used variations on this design. Type 1 (**Fig. 1C i)** and Type 2 (**Fig. 1C ii)** restrict the possible connections to a narrow region ∼75µm from the soma and differ only in the time taken for the presynaptic signal (i.e., input) to propagate to each postsynaptic node (i.e., output). Type 3 (**Fig. 1C iii)** places more submicron channels evenly around the perimeter of each postsynaptic node and provides a larger space for more typical *in vitro* neuron morphology. Each circuit type was implemented in a circuit array (**Fig. 1B i**) designed to be fully compatible with commercially available MEAs so that each node can be aligned with, at least, one microelectrode **(Fig. S1)**.

**Figure 1:**
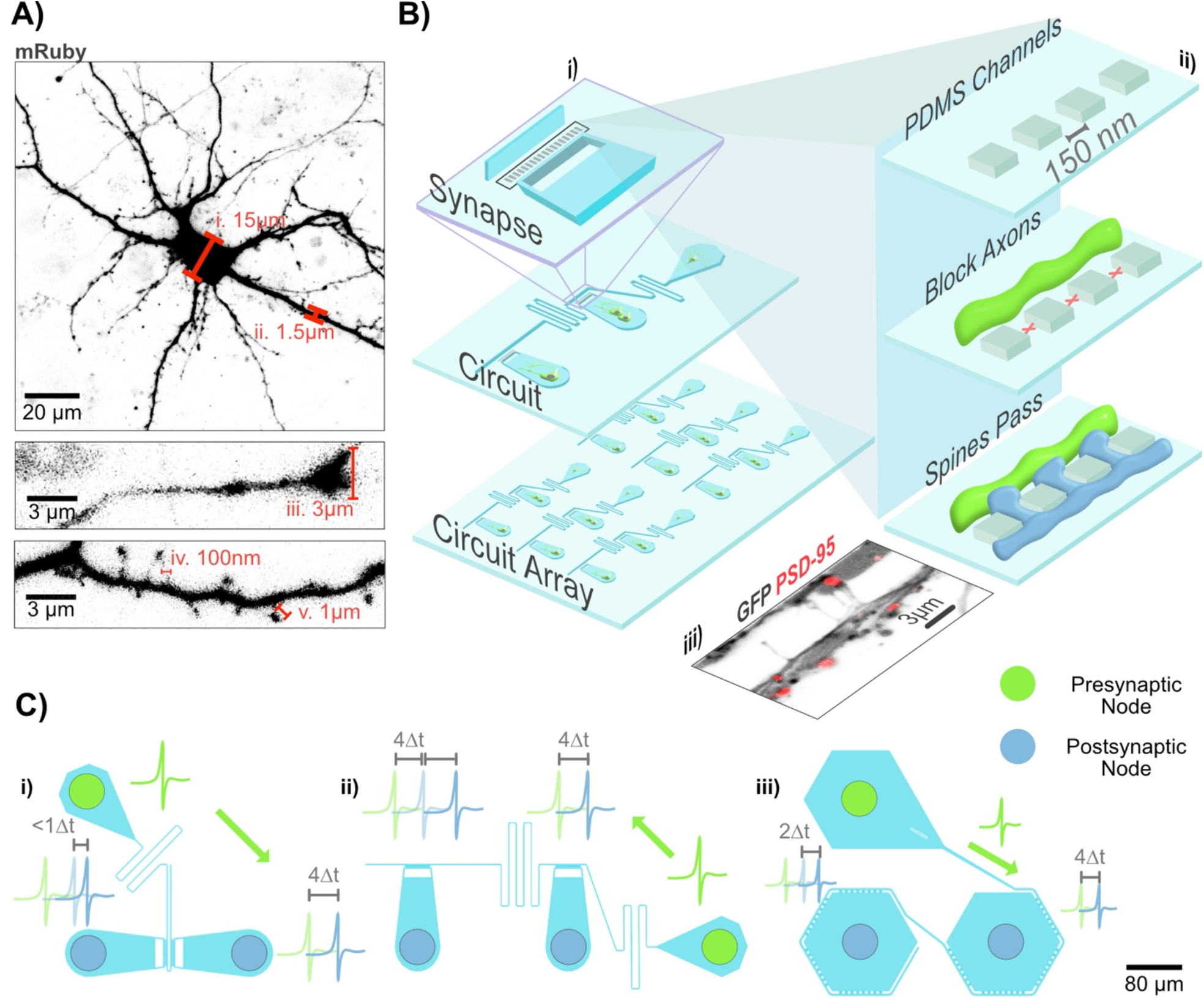
Principle of Multiscale Patterning of Neuron Morphology. **(A)** Neuron expressing mRuby. The text labels indicate typical sizes of relevant subcellular features for rat hippocampal neurons *in vitro*: *i)* the cell body; *ii)* dendrite; *iii)* axon growth cone; *iv)* spine neck; *v)* spine head. **(B)** *i)* Multiscale PDMS devices are used to pattern arrays of circuits with millimeter features and nanochannels for synaptic connections. *ii)* Using feature sizes well below the axon diameter (450nm) but larger than the spine neck width (100nm), synapse formation between the pre and postsynaptic cells can be restricted to the patterned region. *iii)* Fluorescent image of dendritic spines crossing through the nanochannels. Green fluorescent protein (GFP) expressing neurons show spines crossing the nanochannel region. Red fluorescent protein (RFP) tagged postsynaptic density protein 95 (PSD-95) indicates that synapses can form across the channels. **(C)** Neuronal circuit designs consisting of one presynaptic (input) group of neurons with two postsynaptic (output) groups of neurons. The green arrow indicates the intended direction of action potential propagation. i) Type 1 structure where the presynaptic potential should arrive at each postsynaptic node near-simultaneously. ii) Type 2 structure where the presynaptic potential reaches each subsequent postsynaptic node with a significant delay. iii) Type 3 structure with more space for neuron growth and crossing nanochannels around the perimeter of the cell. Circuit designs are to scale, but activity propagation is schematized.

To implement designs utilizing this principle of size restriction, we first needed to develop a fabrication protocol able to reliably generate 150 nanometer features with tolerances on the order of +/-20 nanometers and other features spanning a few millimeters that can be replicated in a biocompatible and transparent material. Due to the large number of replicates typically needed for *in vitro* neuroscience experiments, scalability of the fabrication method was an additional requirement. Soft lithography, the most used method for neuron guidance *in vitro*,^6,7^ has the potential to meet these requirements, but resolution limits in both the master mold and the polymer needed to be addressed. Conventional photolithography using either photomasks or direct laser writing, while able to generate wafer scale molds quickly, is unable to do so reliably at resolutions of 1µm or below **(Fig. S2)**. Creating a mold with directed self-assembly using colloidal lithography or block copolymer lithography would be able to generate nanoscale features reliably at the wafer scale,^38^ but features are typically limited to lines, pillars, tubes, and other regular geometries evenly spaced across the entire wafer when not supported by other patterning techniques.^42^ Alternatively, electron beam lithography (E-beam) can reliably write arbitrary shapes at the wafer scale with sufficient spatial resolution. The main drawbacks of E-beam are that it is a serial process requiring extensive infrastructure making it both slow and expensive to generate dense features at scale. Mix and match lithography, where E-beam lithography and photolithography are used in sequence for different feature layers, addresses this issue by allowing sparse writing of size-critical features with E-beam while larger features can be addressed using standard photolithography techniques.^43^ While this technique has been used in the semiconductor industry, the literature is sparse and to our knowledge has not been used to fabricate molds for soft lithography.

The devices we fabricated required three layers: one for the spine restriction, a second for axon guidance, and a third for the placement of the cell bodies. To generate the nanoscale features required for the spine restriction, a thin layer of SiO2 (∼350nm) was deposited on a Si wafer followed by a thin layer of photoresist that was then patterned using E-beam before the structures were etched in the SiO2 using reactive ion etching (**Fig. 2A i-iv**). After stripping the remaining photoresist, the two subsequent layers were patterned in SU8 using conventional photolithography (**Fig. 2A v-viii**). The major limitation in this process was the alignment tolerance between the SiO2 layer and the first SU8 layer (+/-1.5µm) necessitating an overlap of that amount. With fabrication of the master mold complete, it was cleaned and coated in an antiadhesive silane in preparation for soft lithography. While Sylgard 184 is the most used PDMS, it is not suitable for feature sizes below 500nm.^44^ Instead, a stiffer version of PDMS developed at IBM known as hard PDMS (h-PDMS),^45^ suitable for sub 100nm features at aspect ratios close to 1:1 and sufficient pitch to avoid lateral collapse, was used.^46^ To avoid demolding and handling issues, we used two-layer composite structures following a similar protocol to one in the literature^47^ where a relatively thick layer of a softer PDMS is coated over a partially cured thin layer of h-PDMS (**Fig. 2B**). SEM images of a circuit array, a circuit and spine crossing region for one of the master molds and for the h-PDMS are shown in **Fig. 2C** and **Fig. 2D** respectively. In all designs the channel size was 150nm wide and 300nm long, but different pitches were tested. While a successful mold could be produced for each pitch, faithful replication in h-PDMS was inconsistent when the center-to-center distance was 275nm, the smallest tested, as can be seen in **Fig. 2D vi** where some of the nanochannels are undergoing edge collapse.

**Figure 2:**
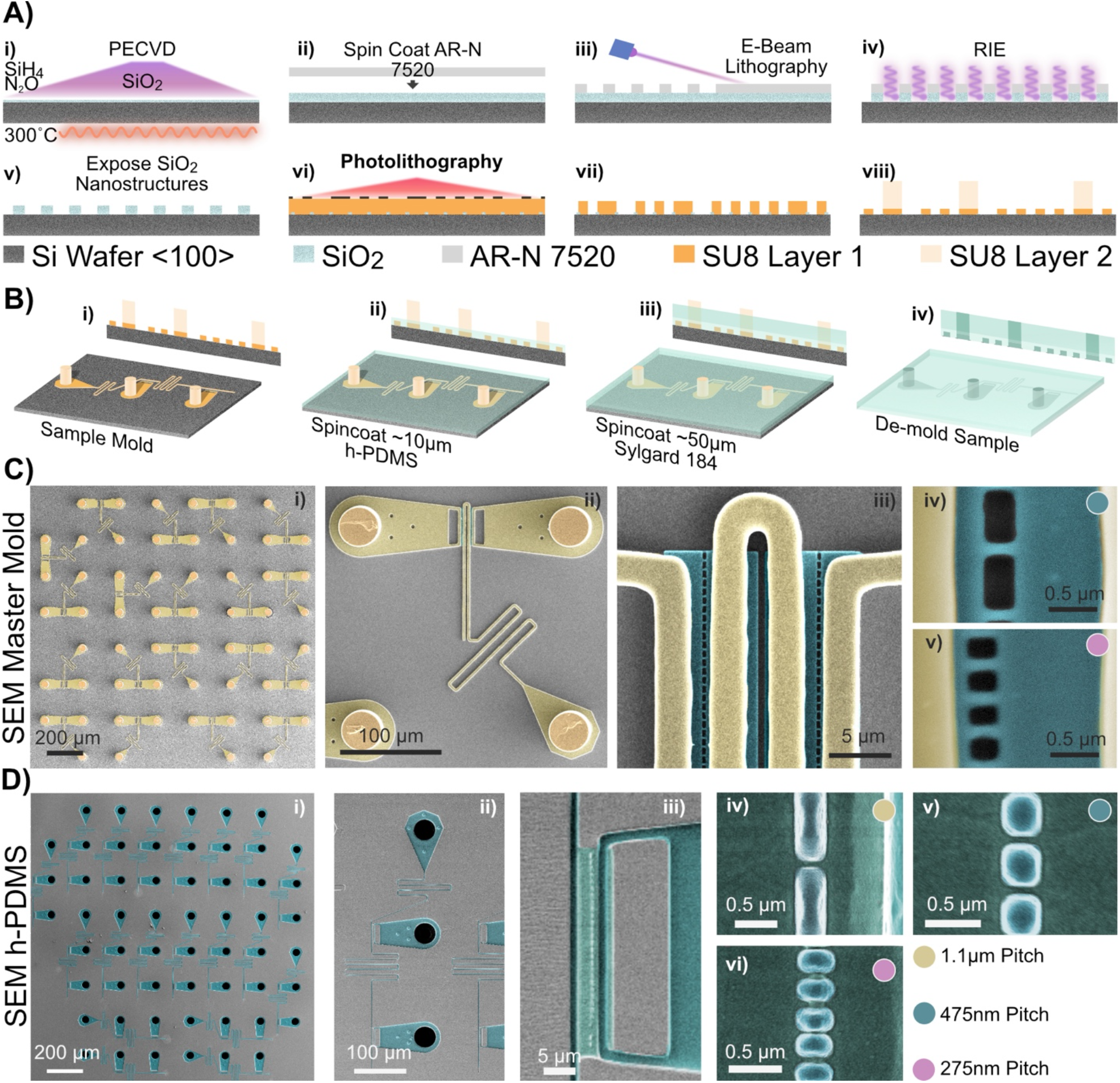
Master Mold and h-PDMS Devices. **(A)** Mix and match lithography process for master mold fabrication: *i)* Plasma Enhanced Vapor Deposition (PECVD) of 360nm of SiO_2_ on a <100> silicon wafer; ii) Spin coat the same thickness of E-beam compatible photoresist; iii) Expose the photoresist and remove uncrosslinked material; iv) Reactive ion etch to remove exposed SiO_2_ with the Si wafer acting as an etch stop; v) Strip the remaining photoresist and clean the sample for further processing; vi) The two subsequent layers are made using conventional photolithography where SU8 is exposed through a photomask; *vii)* The first SU8 layer is 2µm thick and defines the axon guidance channels; *viii)* The second SU8 layer is 75µm thick and defines the cell placement wells. Note: Images are not to scale. **(B)** PDMS structure fabrication: *i)* Cartoon of the master mold; *ii)* Spincoat a thin layer of h-PDMS to cover the first two layers of the mold; *iii)* Spincoat a thicker layer of Sylgard 184 after the h-PDMS partially cures ensuring the pillars for the wells are not covered. *iv)* Gently demold the composite PDMS membrane as high stress can crack or break the h-PDMS. **(C, D)** SEM micrographs of: *i)* Circuit array; *ii)* Individual circuit; *iii)* Spine crossing region; *iv-vi)* 150nm wide channels separated by various distances for the master mold; **(C)** and a PDMS membrane **(D)**. The micrograph for *Di* was taken using a Hitachi SU5000 (2.5 kV), whereas a Magellan 400 FEI was used for the master mold (5.0 kV) and for all other h-PDMS images (2.0kV).

### High-throughput method for circuit isolation

Each PDMS membrane comprises wells (one per node) which neurons fall in and adhere to the substrate after random cell seeding. However, when dealing with thin PDMS membranes (< 0.1 mm thick) the neurons landing on top of the PDMS membrane cannot be neglected; these neurons may connect to those neurons within the nodes and break circuit isolation **(Fig. S3A)**. Alternative methods (without 3D confinement) that precisely position and connect neurons in culture, such as micropatterning of spots and lines,^23,34^ are difficult to parallelize and lack long-term and fine control over circuit connectivity. Thus, for the precise isolation of each circuit, we opted to refine the PDMS nano-microstructures’ substrate preparation and cell culturing protocol (schematized in **(Fig. 3A)**). First, we bonded the nano-microstructures to the glass substrate **(Fig. 3A i-ii)**, which prevented axonal outgrowth under the h-PDMS even at late stages of maturation. To prevent cell deposition and axonal outgrowth on top of the PDMS layer (as in **Fig. S3A**), we aligned a second PDMS membrane (PDMS seeding mask) over the nano-microstructure membrane **(Fig. S3B)**. These seeding masks were comprised of through-holes and acted as a sacrificial layer allowing for selective PDL coating **(Fig. S3C)** of the nano-microchannels and cell deposition in the desired nodes **(Fig. 3A iii-vi)**. A similar approach has also been employed to construct isolated micro 3D cultures connected via microchannels.^35^

**Figure 3:**
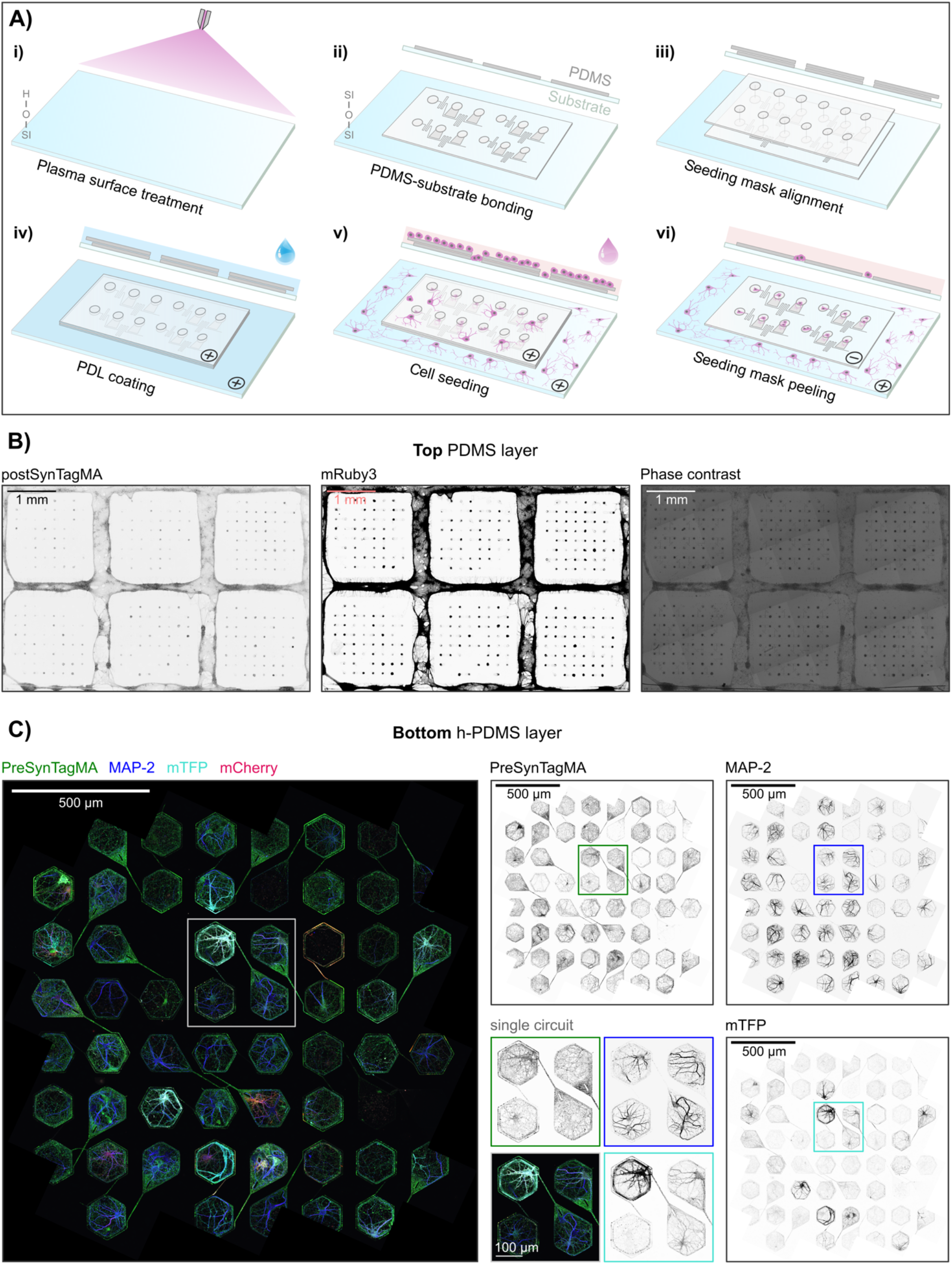
Cell culture method for circuit isolation. **A)** Schematic of the steps needed for substrate preparation and culturing of isolated neuronal circuits: *i)* surface treatment by oxygen plasma cleaning; ii) h-PDMS/PDMS nano-microstructures bonding to the substrate (typically glass); iii) alignment of the PDMS seeding mask through-holes; iv) PDL coating and washing steps; v) replacement with neuronal medium and cell seeding; vi) peeling of the PDMS seeding mask after cell adhesion. A cross-section of each step result is shown on top of the 3D schematic. **(B)** Confocal microscopy mosaic of the top of the PDMS nano-microstructures at 9 days *in vitro*. Neurons expressing postSynTagMA (nuclei and spines) and mRuby3 (whole morphology). **(C)** Confocal microscopy mosaic of the bottom of the h-PDMS nano-microstructures at 20 days *in vitro*. Neurons expressing preSynTagMA (presynaptic terminals), Brainbow (mTFP/mCherry for differential whole-morphology expression) and stained for MAP-2 (dendrites).

At 9 DIV, 98% of the nodes (616 out of 631, from 12 independent circuit arrays) did not establish any connection over the top of the PDMS **(Fig. 3B)**. Thus, the great majority of the inter-node connections were established strictly via the nano-microchannels. **Fig. 3C** shows a representative circuit array at 20 DIV. At 20 DIV, viable neurons could be identified in 93% of the nodes (130 out of 140, from 3 independent circuit arrays), which demonstrates that this cell culture method and circuit design are amenable to the long-term experiments needed for a wide range of applications (e.g., plasticity studies). Moreover, the method is scalable, allowing for the preparation of thousands of circuits (up to 19 circuits per MEA-compatible circuit array) per cell preparation.

In summary, the presented devices and cell culture method allowed for reproducible and high-throughput formation of isolated neuronal circuits. Neurons were deposited in specified areas (i.e., nodes) and expected to extend their processes along the coated nano-microchannels thereby forming microcircuits. Although we could not precisely control the number of neurons per node, with the employed cell seeding densities it typically did not exceed just a few (< 5 neurons per node). In the future, a precise number of neurons per node may be achieved via, for example, controlled single-cell deposition using the FluidFM system.^48^

### Multiscale subcellular neuronal patterning

We tested if the proposed circuit designs imposed the multiscale subcellular patterning needed for the consistent formation of feed-forward neuronal circuits. For this to occur, it was essential that different rules of restriction were followed: 1) presynaptic node’s neurons cannot extend dendrites to the postsynaptic nodes; 2) axons cannot cross the nanochannels (so that postsynaptic node’s neurons do not extend axons to other nodes); 3) dendritic spines can cross the nanochannels to form synapses. We successfully applied these rules via a combination of length- and size-dependent physical restrictions that comply with the physiological neuronal dimensions **(Fig. 1)**.

As expected from the literature,^26^ dendrites did not grow for more than 200-300 µm, thus the presynaptic node’s long emitting microchannel isolated axons in a length-dependent manner **(Fig. 4A)**. However, the extending axons did not cross the nanochannels due to their larger size. Thus, axons from the presynaptic node’s neurons did not enter the postsynaptic node (presynaptic axon restriction) **(Fig. 4B** and **Fig. S4)**. At the same time, axons originating from the postsynaptic node did not enter the afferent microchannel (postsynaptic axon restriction) **(Fig. 4C)**. Finally, the nanoscale dimensions ensured feed-forward connectivity between the circuit nodes, with only dendritic spines being able to cross the nanochannels. These dendritic spines may encounter axons emitting from the presynaptic node and form putative synapses, as indicated by the presence of the synaptic scaffolding protein PSD-95 **(Fig. 4D)**.

**Figure 4:**
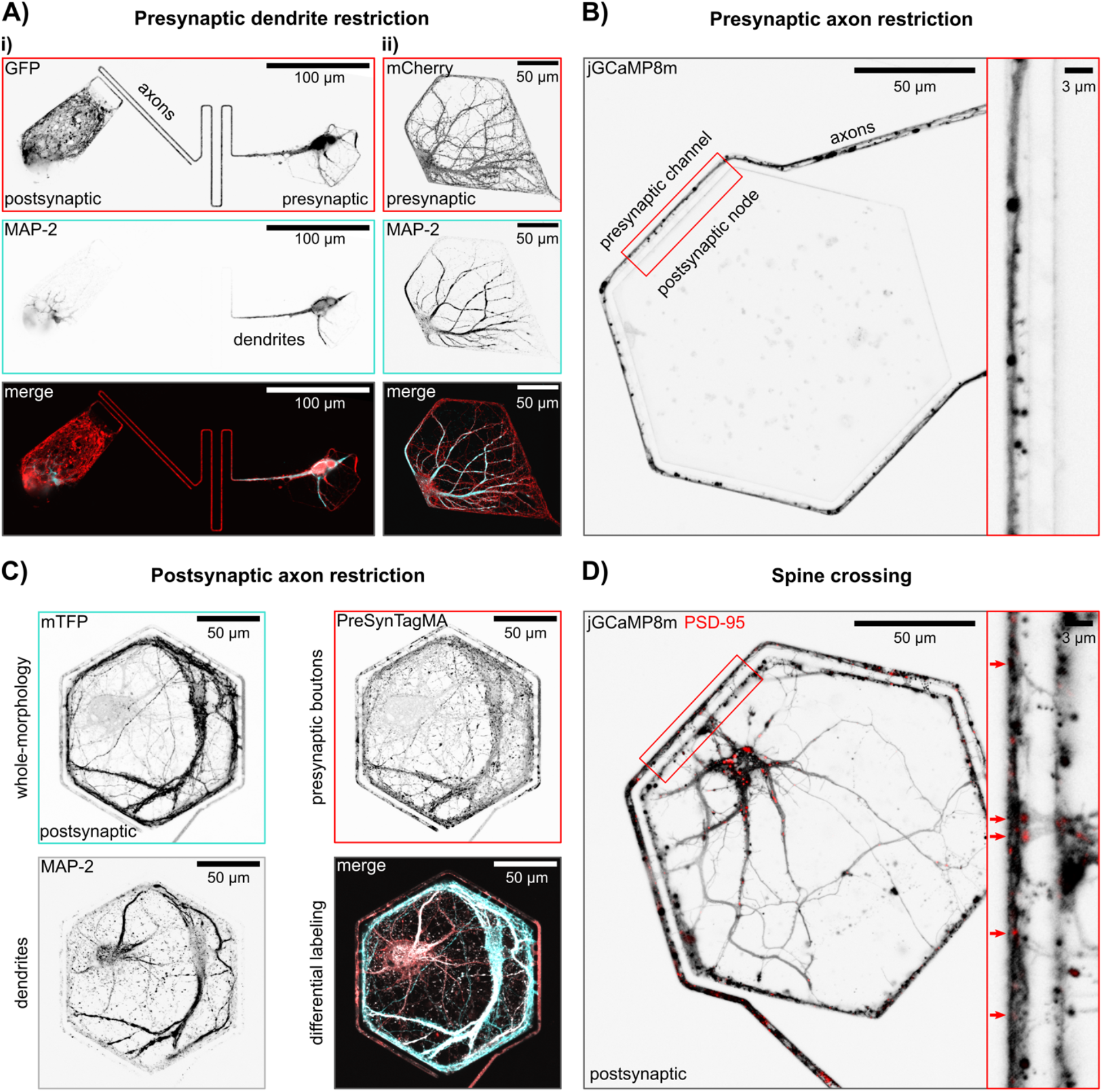
Multiscale subcellular neuronal patterning. **A)** Examples of presynaptic dendritic outgrowth restriction. In both, the long microchannel emitting from the presynaptic node prevents dendritic outgrowth to the postsynaptic nodes. i) Dendrites enter the emitting microchannel in Type 1-2 node-designs due to lack of space in the presynaptic node, but they cannot reach the postsynaptic nodes. ii) Type 3 node-design provides a larger space for more typical *in vitro* neuron morphology; thus, dendrites arborize within the node. **B)** Example of presynaptic axon restriction. The nanochannels restrict axonal outgrowth from the presynaptic node to the long microchannel, preventing invasion into the postsynaptic nodes. The respective intact nanochannels can be seen in **Fig. S4. C)** Example of postsynaptic axon restriction. The nanochannels prevent axonal outgrowth from the postsynaptic nodes into the long microchannel. Note that the differential protein expression (via a Brainbow construct) allows for morphological separation of presynaptic (expressing PreSynTagMA) and postsynaptic (expressing mTFP and PreSynTagMA) neurons. **D)** Example of spine crossing and putative synapse formation. The nanochannels allow for dendritic spine crossing, thus synapse formation between the pre- and postsynaptic nodes (arrows indicate putative synapses).

This device concept positions synapse formation in a controlled area that can facilitate the high-throughput analysis of synaptic features (e.g., synapse structure, number, or function) in-between connected nodes. Since differential labelling strategies (e.g., Brainbow constructs) can be used to differentiate processes originating from the pre- or postsynaptic nodes **(Fig. 4C)**, the separation of overlapping processes (e.g., axons/spines) could be achieved. A typical *in vivo* approach is to use sparse labelling methods, which helps to distinguish and reconstruct a given neuron’s morphology within the neuropil, but do not label most connecting neurons. Our bottom-up approach allows for labeling all neurons within the microcircuit, as well as their inter-node connections, paving the way for continuous mapping of the connectome and synaptome of neuronal microcircuits. Previous *in vitro* studies have compartmentalized synapse formation to desired areas via microcontact printing^49,50,51^ or tripartite microfluidic chambers.^52^ These approaches can be used effectively to exclude neuronal somata and promote randomized synapse formation in specified large areas. However, they do not allow for precise engineering of neuronal microcircuits (e.g., with feed-forward connectivity), nor the straightforward identification of which neurons (or nodes) are connected.

### Functional synaptic connections in-between nodes

Neuronal electrical activity reliably leads to calcium influx; thus, calcium imaging has long been used as a proxy for neuronal activation. As such, calcium signals in axons and dendritic spines are used as proxies for presynaptic and postsynaptic activity, respectively. Since these small subcellular compartments are difficult to probe with electrophysiological techniques, the localized calcium transient is often used as a readout of synaptic activation.^53^ However, both *in vivo* and *in vitro*, the study of synaptic calcium signaling is made difficult by the intricate neuronal network, with a high-density packing of dendritic spines and intercrossing axonal arbors **(Fig. S5)**. Here, we took advantage of the synaptic compartmentalization in a well-defined area and, as a proof-of-concept, performed calcium imaging to explore if the circuit nodes established functional synaptic connections. This synaptic compartmentalization allowed for the straightforward identification of putative synapses and the assessment of their activity (active or silent) **(Fig. 5)**.

**Figure 5:**
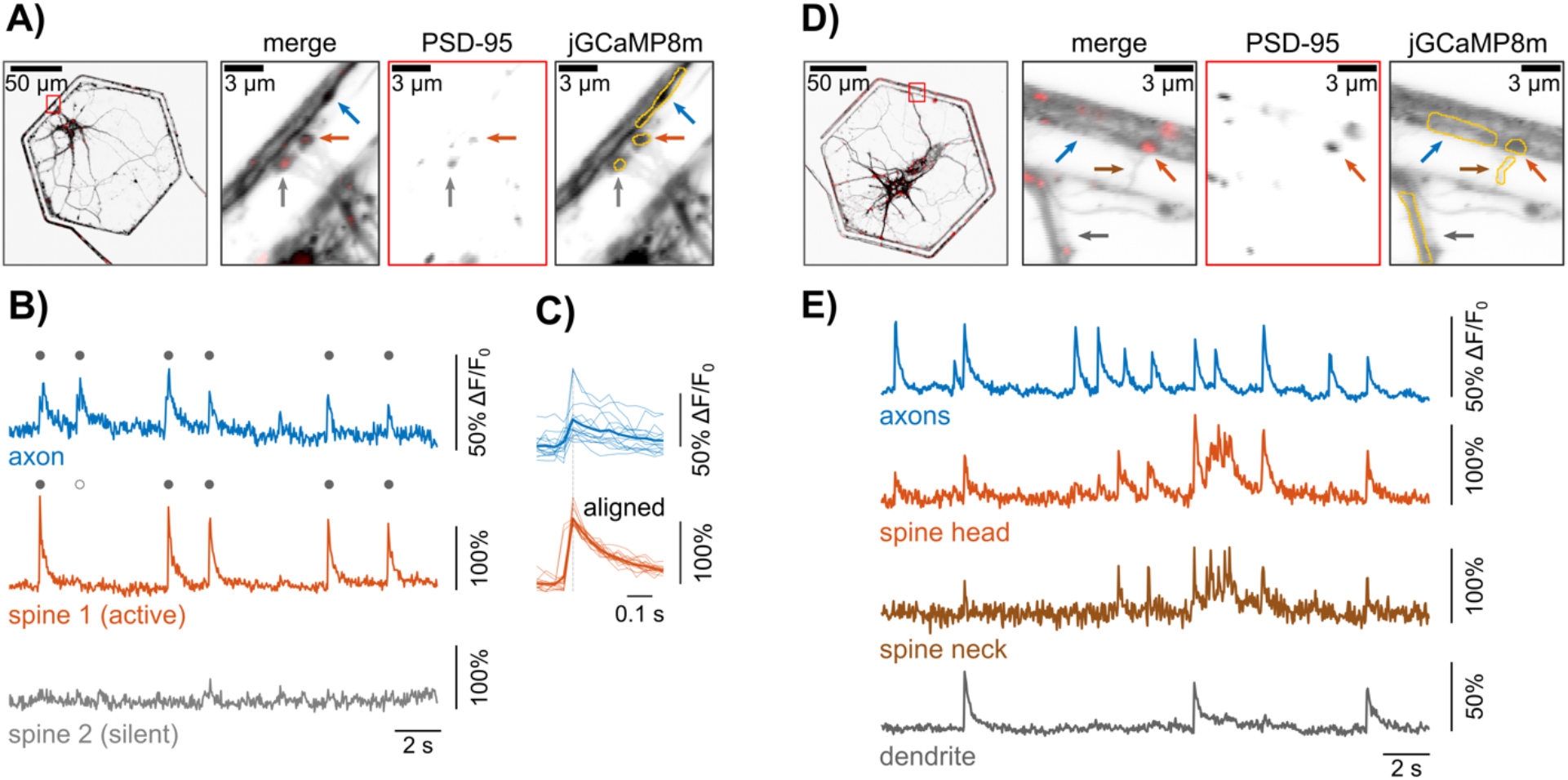
Circuit nodes establish functional synaptic connections. **A)** Example of circuit neuron expressing PSD-95 and jGCaMP8m. Zoom-ins of the red inset with two identifiable spines and an emitting axon. Regions of interest (ROIs) used for the analysis are delineated in the jGCaMP8m channel. **B)** Calcium traces of the axon and spines’ activity. Filled dots represent coincident axon and spine calcium events. **C)** Event coincidence of spine 1 and axon shown by aligned peaks of all spine events (n = 15) and corresponding axon traces (average traces are shown as thick lines). **D)** Example of circuit neurons expressing PSD-95 and jGCaMP8m. Zoom-ins of the red inset with one identifiable spine (head and neck) and respective dendritic shaft. ROIs used for the analysis are delineated in the jGCaMP8m channel. **E)** Calcium traces of the multiple sub compartments.

Neurons expressing jGCaMP8m at 14 DIV were used in the calcium imaging experiments. The high SNR of this new sensor^54^ allowed for the morphological identification of single axons within the emitting microchannel, as well as single dendritic spines. We could readily identify potential synapses via PSD-95 labeling **(Fig. 5A)**. To assess synaptic function, we performed paired recordings of spontaneous pre- and postsynaptic activity. In an example axon–spine pair (i.e., putative synapse), all the postsynaptic events in the dendritic spine were coincident (i.e., within ∼33 ms; the duration of one frame) with presynaptic input **(Fig. 5 B-C)**. Even though not all presynaptic events induced postsynaptic events **(Fig. 5B)**, as is expected for functional synapses, the local calcium signals from the dendritic spine head were highly correlated with calcium signals from the apposing axon (Pearson correlation, *r* = 0.54) **(Fig. 5 B)**. Moreover, with the achieved level of compartmentalization, we could analyze the full spatiotemporal pattern of calcium signaling along multiple neuronal compartments, as shown in **Fig. 5 D-E**. Currently, most calcium imaging experiments focus in either the somatic, axonal, dendritic or spine compartment due to the spatial scaling difficulties and the processes signal overlap.^53^ Here, we could discriminate and record calcium signals from all these compartments simultaneously, with a single sensor **(Fig. 5E)**, and without the need to stimulate another compartment externally (e.g., via current injection). In the future, dual-color imaging of the pre- and postsynaptic compartments (e.g., with red and yellow XCaMPs) can further facilitate the compartments discrimination.^55^ Ultimately, this patterning method may provide a unique tool for the study of synaptic function and mechanisms associated with synaptic plasticity that transverse multiple compartments, such as backpropagating action potentials.

Recent endeavors have used electron microscopy to map 1 mm^3^ of preserved human cerebral cortex (the equivalent of ∼1-pixel in a functional magnetic resonance imaging (fMRI) scan), at a nanometer scale that is required to identify individual synapses. The resulting 1.4-petabyte volume of raw data could only be scrutinized via automated procedures, which revealed 133.7 million putative synapses.^56^ Despite the important insights that this technical approach (i.e., ultrastructural studies) may reveal on the fundamentals of synaptic organization and variability, it only provides a snapshot of the synaptome with little to no information of synaptic strength or plasticity. Classically, *in vitro* patch-clamp electrophysiology, despite its invasiveness, has been the go-to tool for the study of synaptic function and plasticity in short-term experiments.^5,57^ Recently, patch-clamp electrophysiology, followed by electron microscopy reconstruction of all the putative synapses between the recorded neurons, revealed a linear correlation between synapse size and strength.^58^ This technique combination ought to reveal more insights into the structure/function relationship of synapses, but its’ technical complexity, low-throughput, and incompatibility with long-term experiments (i.e., more than a few hours) preclude the mapping and following of synaptic connectivity during prolonged physiological and/or pathological states. We envision our proposed device and concept can help fill this technological gap, by providing a testbed for long-term enquiry of circuit and synaptic function.

## CONCLUSION

We created a device and methods to establish multiscale, well-defined, node-based microcircuits *in vitro*. Node-based circuits with various degrees of intra- and inter-connectivity, such as the ones established in this work, are interesting not only in their own right but also since *in vivo* and *in silico* studies support the hypothesis that information processing occurs in node-like circuit topologies.^14^ By reducing the circuit complexity this novel device may be a valuable tool for the study of fundamental concepts in neuroscience, such as information storing and processing.

We demonstrated that these microcircuits form functional synaptic connections, thus the opportunities for the short- and long-term study of synaptic function are nearly endless. Importantly, the thin nano-microstructure membranes (∼60 µm) are compatible with live high-resolution microscopy, thus future studies may take advantage of an extended and increasing repertoire of compatible optical (e.g., optogenetics) and electrophysiological technologies (e.g., MEAs or patch-clamp) to study and manipulate the circuit connectivity. We anticipate that this approach to multiscale patterning in combination with non-invasive functional sensors, such as calcium imaging or MEAs, can be used to study the fundamental characteristics of neuronal networks in long-term experiment opening several new experimental opportunities with relevance not only to the burgeoning field of bottom-up neuroscience but also top-down experimental and computational neuroscience.

## METHODS

### Mold design and fabrication

Fabrication of the master mold (Wunderlichips) was done using a combination of E-Beam lithography, reactive ion etching (RIE) and photolithography. Full details on the parameters used in each fabrication step and the design rules can be found in the **supplemental methods**. Design of the structures was done using AutoCAD 2020 (AutoDesk) with 525 circuit arrays fitting on a 4-inch wafer. A <100> silicon wafer (Silicon Materials) was cleaned in oxygen plasma (PVA Tepla, GIGAbatch 310M) and baked at 200°C to clean the surface. Next plasma enhanced chemical vapor deposition (Oxford Plasmalab 100 PECVD) was used to deposit 360nm of SiO2 **(Fig. 2 A i)**. Thickness of the oxide layer was measured with an ellipsometer (Bruker Dektak-XT) after deposition. The wafer was again plasma cleaned and baked before an estimated 360nm layer of AR-N 7520 photoresist was spin coated at 500 rpm for 5s followed by 6000rpm for 60s **(Fig. 2 A ii)** and the desired features exposed via by E-beam (Vistec EBPG 5200+) **(Fig. 2 A iii)**, and the resist developed in a mixture of AR300-47 and water (4:1) for 60 seconds. The exposed SiO_2_ was removed using RIE (Oxford NGP 80) with a 50:1 mixture of CHF3 and O2 for 24 minutes **(Fig. 2 A iv)**. As the SiO2 features were not ideal for subsequent alignment steps, AZ6612 was spin coated on the wafer (40s, 4000rpm), the alignment marks exposed, and a higher aspect ratio Si alignment mark exposed using an isotropic Si reactive ion etch. The remaining photoresist layers were then stripped, the surfaces were cleaned with oxygen plasma, and the SiO2 structures were visually and mechanically inspected (**Fig. 2 A v**). Two chromium photomasks were fabricated with a direct laser writer (Heidelberg DWL 66+) and used in the subsequent photolithography steps. Prior to spin coating the wafer was once again cleaned by oxygen plasma, then approximately 2µm of SU8 2002 (Kayaku) was spin coated on the wafer (500 rpm for 5s, 3000 rpm for 30s). It was soft baked and the resist was mechanically removed from the alignment marks and was left to rest overnight. The first photomask and wafer were then aligned in a mask aligner (Süss MA6) using vacuum contact and the photoresist was exposed with pulsed 80mJ/cm^2^ UV light (**Fig. 2A vi, vii**). After a post exposure bake, a 75µm thick layer of SU8 3050 was spin coated on the wafer (500 rpm for 10s, 1700 rpm for 30s) and soft baked before alignment with the second photomask under hard contact. The photoresist was then exposed with 300mJ/cm^2^ pulsed UV light, baked, developed in mrDEV 600, rinsed with IPA, dried, and fully crosslinked under flood exposure of UV light **(Fig. 2A viii)**. The mold was then hard baked at 200°C for 900 seconds before visual and mechanical inspection.

### Nano-microstructure fabrication

Prior to soft lithography, a layer of perfluoro-octyl trichlorosilane (448931 Sigma Aldrich) was chemically vapor deposited on the surface of the master mold to facilitate demolding of the PDMS membrane. As the most used type of PDMS, Sylgard 184 (Dow Corning), is unsuitable for reliably reproducing sub micrometer features^59^ an approximately 10µm layer of h-PDMS, a stiffer PDMS capable of replicating sub 100nm features,^47^ was first deposited on the mold to ensure faithful reproduction of all micron/submicron features **(Fig. 2B ii)**. A 50µm thick layer of Sylgard 184 was then spun on top **(Fig. 2B iii)** for ease of handling as h-PDMS is comparatively brittle and breaks easily. The h-PDMS (Gelest, PP2-RG07) was mixed in a 1:1 ratio of Part A to Part B as per the manufacturer’s specifications, degassed, spin coated at 3000 rpm for 300 seconds, and partially cured for 30 minutes at 60°C; after which Sylgard 184 (1:10 ratio of prepolymer to crosslinker) was spin coated at 1000rpm for 120 seconds before being degassed and cured at 80°C overnight. Finally, the composite PDMS membrane was gently peeled from the master mold, diced, and stored in air until use **(Fig. 2B iv)**.

### Scanning electron microscopy (SEM)

The PDMS structures were placed on a carbon tape covered SEM stub and connected to the carbon tape with silver paste and dried overnight. Then a 10 nm thick layer of platinum was sputtered on the samples with a CCU-010 Metal Sputter Coater (Safematic GmbH). High resolution images were taken with a Magellan 400 FEI SEM at 2.0 kV at high vacuum unless otherwise specified in which case they were imaged with a Hitachi SU5000 at 2.5kV. The images for each structure were taken at different magnifications. The master mold was coated with 5nm of gold and imaged at 5.0 kV at high vacuum.

### Substrate preparation and surface functionalization

Glass-bottom WillCo dishes (GWST-3522, 22 mm, WillCo Wells, Netherlands) or microelectrode arrays (MEAs) (60ThinMEA200/30iR-ITO-gr, MultiChannel Systems, MCS, Germany) were used as cell substrates. All substrates were cleaned and washed by rinsing with ethanol, 1% SDS and ultrapure water (18MΩ/cm Milli-Q, Merck-MilliPore), followed by blow-drying with N_2_. h-PDMS/PDMS nano-microstructures and seeding masks were individually cut from the membrane using a surgical blade and placed channel side up on clean WillCo dishes for plasma cleaning. A schematic diagram of the device preparation is shown in **Fig. 3A**. For optimal adhesion, contacting surfaces were oxygen-plasma cleaned for 2 min (18W PDC-32G; Harrick Plasma, USA) right before alignment. Then, the nano-microstructures were wetted with drops of isopropanol for facilitating the alignment over the desired substrate. Aligned devices were subsequently placed in a desiccator chamber and vacuum was applied for bonding. Next, seeding masks were aligned over the nano-microstructures using the same methods as above. All alignment steps were performed manually under a stereomicroscope using anti-static tweezers. The resulting devices were again plasma-cleaned for 2 min right before coating with 25 µg/ml poly-D-lysine (PDL, P6407 Sigma Aldrich). After >30 min, PDL was washed out with ultrapure water at least 3 times, each time placing the mounted devices in a desiccator chamber with slight vacuum. Finally, the washing solution was replaced with neuronal medium, and the mounted devices were placed in a cell culture incubator (37ºC, 5% CO_2_) until cell seeding.

### Cell culture

All animal experiments were approved by the Cantonal Veterinary Office Zurich. Primary embryonic rat hippocampal or cortical neurons were isolated from Sprague Dawley embryo rats (E18). Tissues were enzymatically digested in 0.5 mg/ml papain (P4762, Sigma-Aldrich, Switzerland) and 0.01 mg/mL Deoxyribonuclease (D5025-15KU, Sigma-Aldrich) in PBS supplemented with 0.5 mg/mL Bovine Serum Albumin (BSA, 11020021, Gibco, Thermo Fisher Scientific) and 10 mM D-(+)-glucose (G5400, Sigma-Aldrich) for 30 min at 37 °C. Subsequently, tissue fragments were washed once with 10% (v/v) heat-inactivated fetal bovine serum (FBS, F2442 Sigma Aldrich) in Neurobasal Plus medium (A3582901, Thermo Fisher Scientific), and twice with Neurobasal Plus medium. Tissue fragments were then mechanically dissociated with a 5 ml serological pipette and filtered with a 40 µm strainer (CSS013040, Biofil) to exclude remaining tissue clumps. Viable cells were counted using the trypan blue exclusion assay. Around 250k and 180k viable cells were seeded on the prepared WillCo and MEA devices, respectively. After seeding, cells were resuspended thrice to ensure an even surface distribution and entrance on the microwells. Cells were cultured in Neurobasal Plus medium supplemented with 2% B-27 Plus (A3582801), 2% GlutaMAX (61965-026) and 1% penicillin/streptomycin (15140-148; all from Thermo Fisher Scientific) and kept in a humidified incubator at 37 °C supplied with 5% CO_2_. At 1-2 DIV, seeding masks were carefully peeled using sterile tweezers and the culture medium was fully replaced. Half-medium changes were performed every 3-4 days for the remaining time in culture.

### Viral Transductions

Transductions with adeno-associated viruses (AAVs) were performed to generate various morphological labels (e.g., GFP) or sensors (e.g., jGCaMP8m) in the neuronal cultures. Targeted viral load was on the order of ten thousand viral particles per cell for all AAVs. All viral vectors were produced by the Viral Vector Facility (VVF) of the Neuroscience Center Zurich unless otherwise specified (Zentrum für Neurowissenschaften Zürich, ZNZ, Switzerland). A list of all AAVs used in the study is presented in Table S1.

**Supplementary Table 1.**
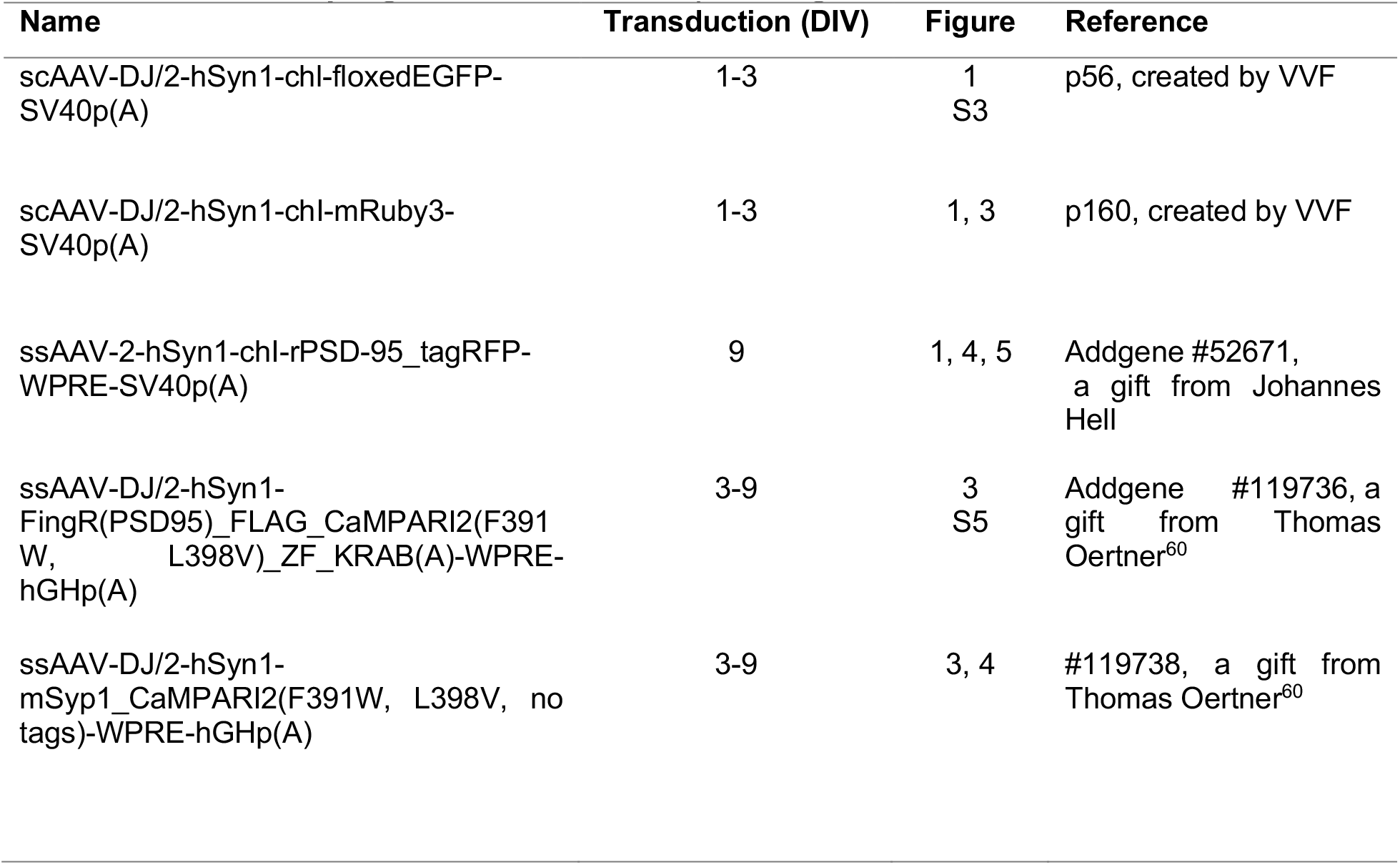
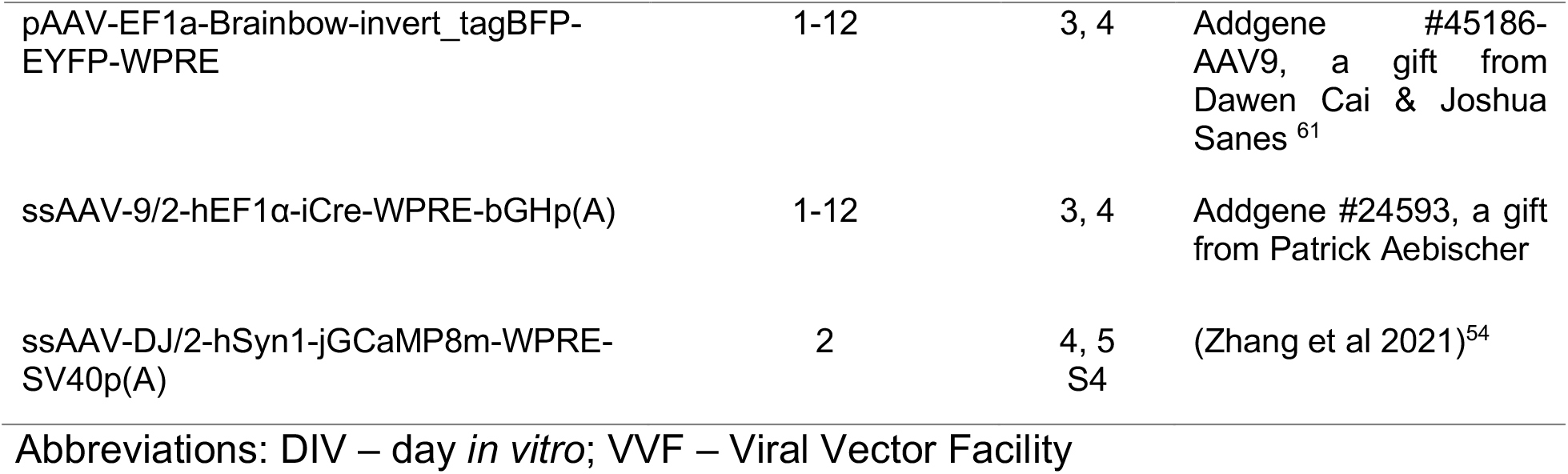
List of all adeno-associated viruses (AAVs) used in the study, their transduction day, figure in which the expressing neurons are shown and references.

### Immunolabeling

Our immunocytochemistry protocol was derived from that in Taylor et al. 2005 for neurons in PDMS structures (Taylor 2005). Briefly, cells were fixed with 4% paraformaldehyde for 30 minutes at room temperature (RT), permeabilized with 0.2% Triton-X for 30 minutes at RT, and blocked with 3% BSA for 2hrs at 37°C. Subsequently, the primary antibody was added to a 3% BSA, 0.2% Triton X solution and left to incubate at 4°C overnight. After three washing steps with PBS, a secondary antibody was added and incubated for, at least, 1h RT. We used a MAP-2 polyclonal antibody (PA5-17646 Invitrogen, 1:1000 dilution) to stain for dendrites, as the primary antibody. The Hoechst dye 33342 (62249, Thermo Fischer Scientific, 1:500 dilution) was used to stain for nuclei.

### Fluorescent Imaging

All fluorescence microscopy was done using a FluoView 3000 (Olympus) confocal laser scanning microscope equipped with temperature and CO_2_ control to maintain a 37°C and 5% CO_2_ environment during live cell imaging (Pecon). Coverslips with a thickness of 170µm were used in all imaging experiments. Physically based z-drift compensation (TruFocus, Olympus) was used throughout experiments to maintain a consistent focal plane during multiarea scans (e.g., Fig. 3A). Laser wavelengths used were 405nm, 445nm, 488nm, 561nm, 594nm, and 640nm. A 60x (UPLSAP060XS2, Olympus) or 30x (UPLSAP030XS, Olympus) silicon immersion objective was used for all high-resolution images while a 10x (UPLFLN10XPH) or 20x (UPLFLN20XPH) air objective were used for coarse imaging. Imaging was done between DIV9 and DIV20 for investigating structure while calcium imaging was done on DIV14 (12 days post transduction).

### Image processing and analysis

Image processing and analyses were performed with ImageJ and MATLAB. For calcium imaging analysis, photobleaching was corrected with an exponential fit to the decay curve. Then, videos were median-filtered (1-pixel radius) before calculating ΔF/F_0_. The median projection (background intensity map) was used to define F_0_. Regions of interest (ROIs) were delineated manually, and ΔF/F_0_ traces were calculated for the subcellular structures of interest. ROIs of axons and dendritic spines were delineated based on typical morphological characteristics (e.g., axon as a thin aspiny process), location (e.g., axons within the emitting microchannel) and labeling (e.g., dendritic spines with PSD-95 puncta).

## Author Contributions

S.W. and J.C.M. designed the study, performed the experiments, and analyzed the data; D.vS. developed the process and fabricated the PDMS devices; A.R. and J.H conducted SEM imaging;. J.C.M and S.W. wrote the manuscript; J.V. and P.A. supervised the study; S.W., J.C.M, P.A., J.V. revised the manuscript; All authors have contributed to the writing and approved the manuscript.

## Funding Sources

J.C.M. was supported by FCT (PD/BD/135491/2018) in the scope of the BiotechHealth PhD Program (Doctoral Program on Cellular and Molecular Biotechnology Applied to Health Sciences). ETH Zurich, the Swiss National Science Foundation, and the OPO Foundation are also acknowledged for financial support.

## Notes

D.vS. operates a company, Wunderlichips GmbH, offering soft lithography services. All other authors have no competing financial interests.

## ACKNOWLEDGMENTS

All the members of the Neuroengineering and Computational Neuroscience (NCN) group (i3S – Instituto de Investigação e Inovação em Saúde da Universidade do Porto) for critical discussions.

