## Supplemental material for "Nanoscale patterning of *in vitro* neuronal circuits"

### Supplementary Information

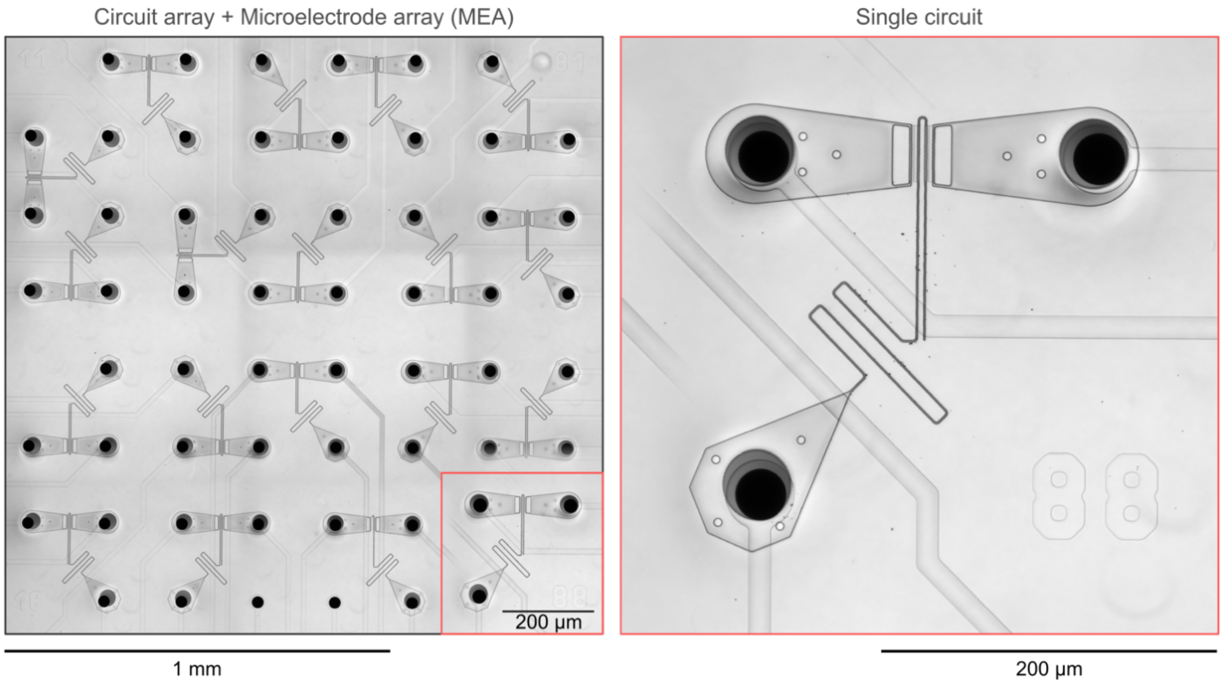


**Figure S1: Combination of a circuit array and a microelectrode array (MEA).** Example light microscopy mosaic image of a circuit array aligned over a commercially available microelectrode array (MEA) (60ThinMEA200/30iR-ITO) composed of 59 recording microelectrodes with 200 µm inter-electrode spacing. Each node encompasses one microelectrode. A zoom-in of a single circuit is shown on the right.

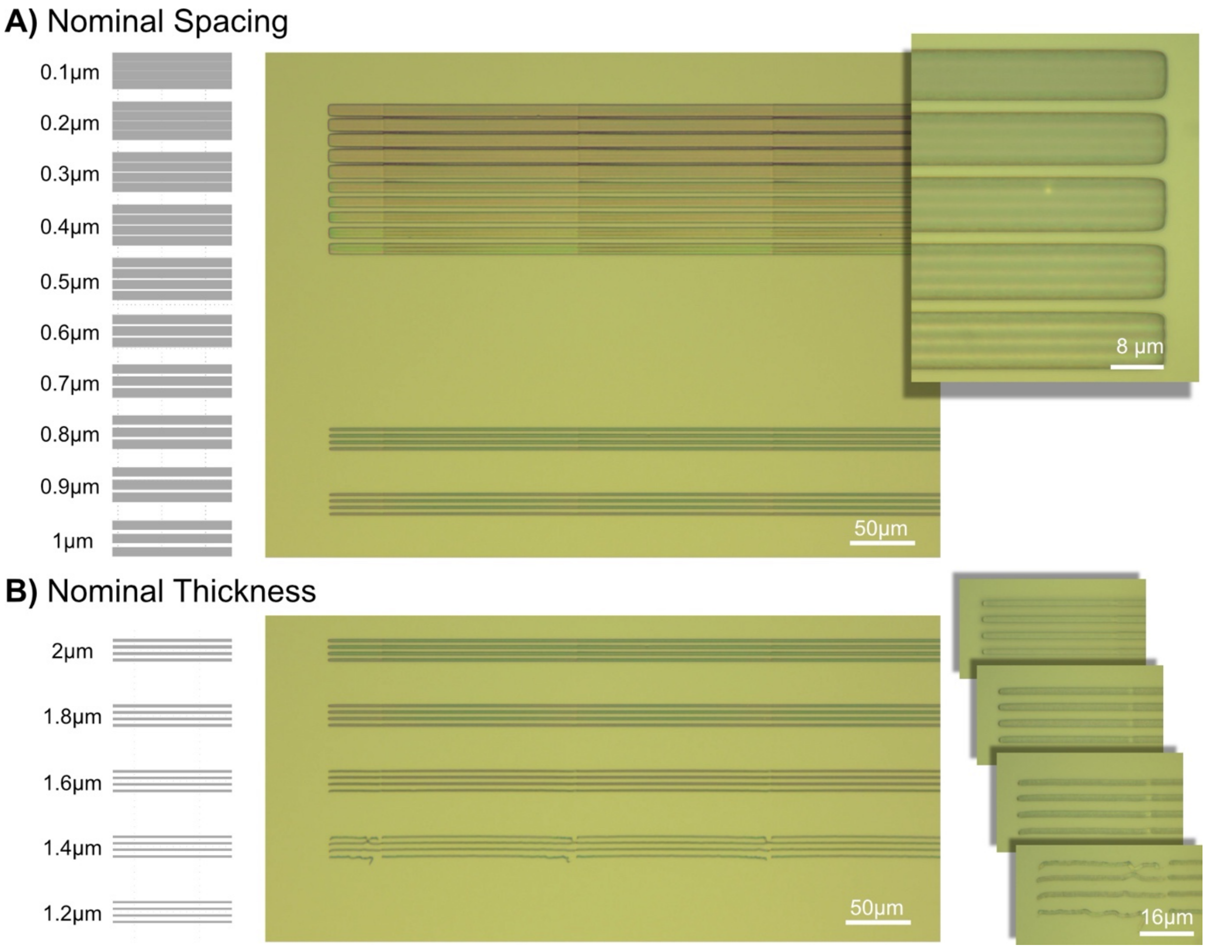


**Figure S2: Resolution test of SU8 2000 for line width and spacing A)** Spacing test of 2µm thick lines, ranging from 0.1µm inter-distance to 1µm inter-distance. The results showed that spacing of 1µm or less are not feasible. **B)** Line width test starting at 2µm and decreasing by 0.2µm each step. Line widths of 1.4µm are possible but have adhesion problems. Line widths below 1.4µm could not be written

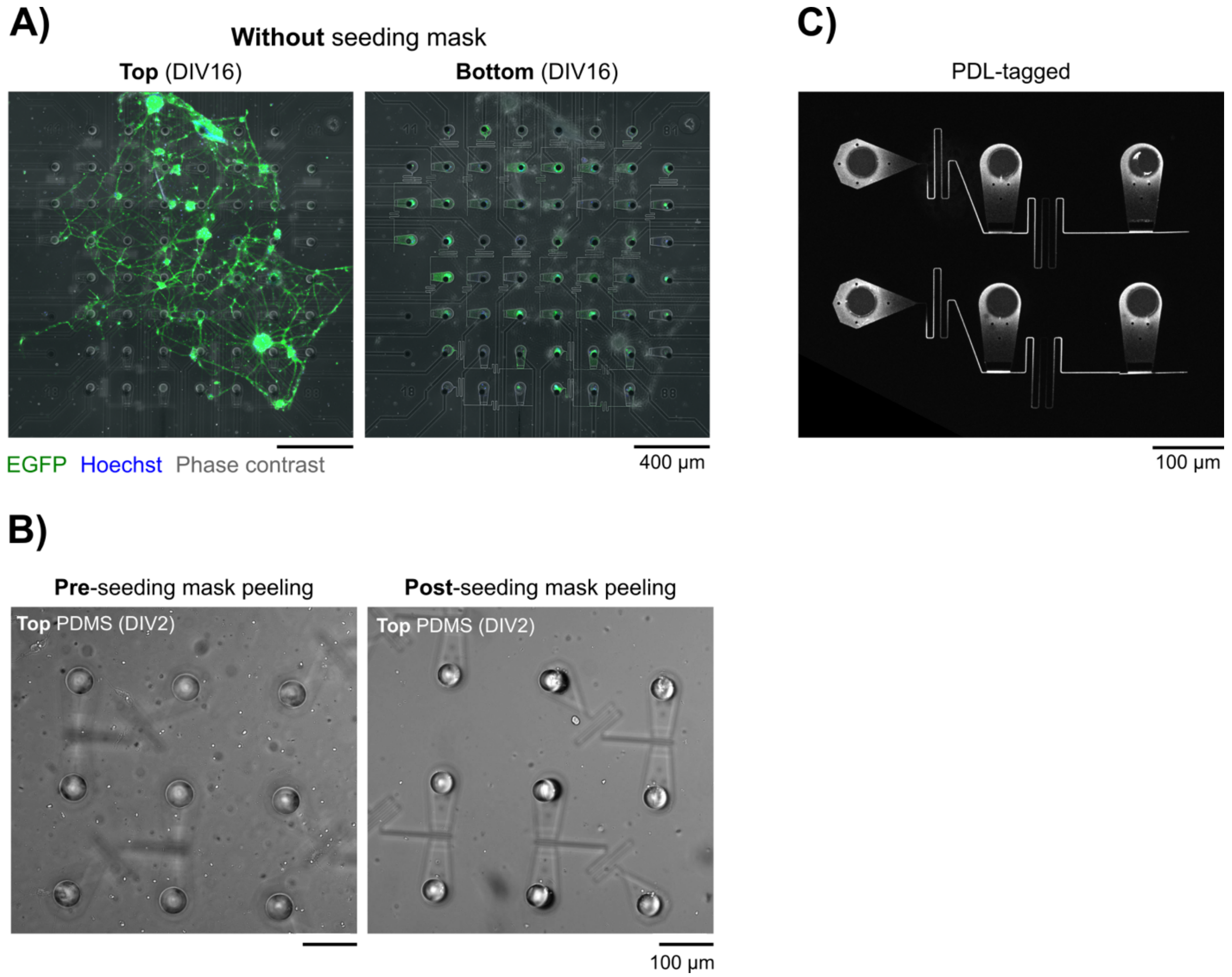


**Figure S3: Seeding masks allow for circuit isolation. A)** Confocal microscopy images of the top and bottom of a nano-microstructures' circuit array which underwent cell culture without a seeding mask. The circuit array is placed on a microelectrode array (MEA). An interconnected neuron network forms on top of the PDMS nano-microstructures. **B)** Example phase contrast images before and after the peeling of the PDMS seeding mask. A neuron-free top surface is obtained. **C)** Confocal microscopy image of mounted nano-microstructures after coating with PDL tagged with a secondary antibody (Alexa Fluor 488). Imaging was performed 24 hours post-coating. The device mounting procedure allows for selective PDL coating of the nano-microchannels.

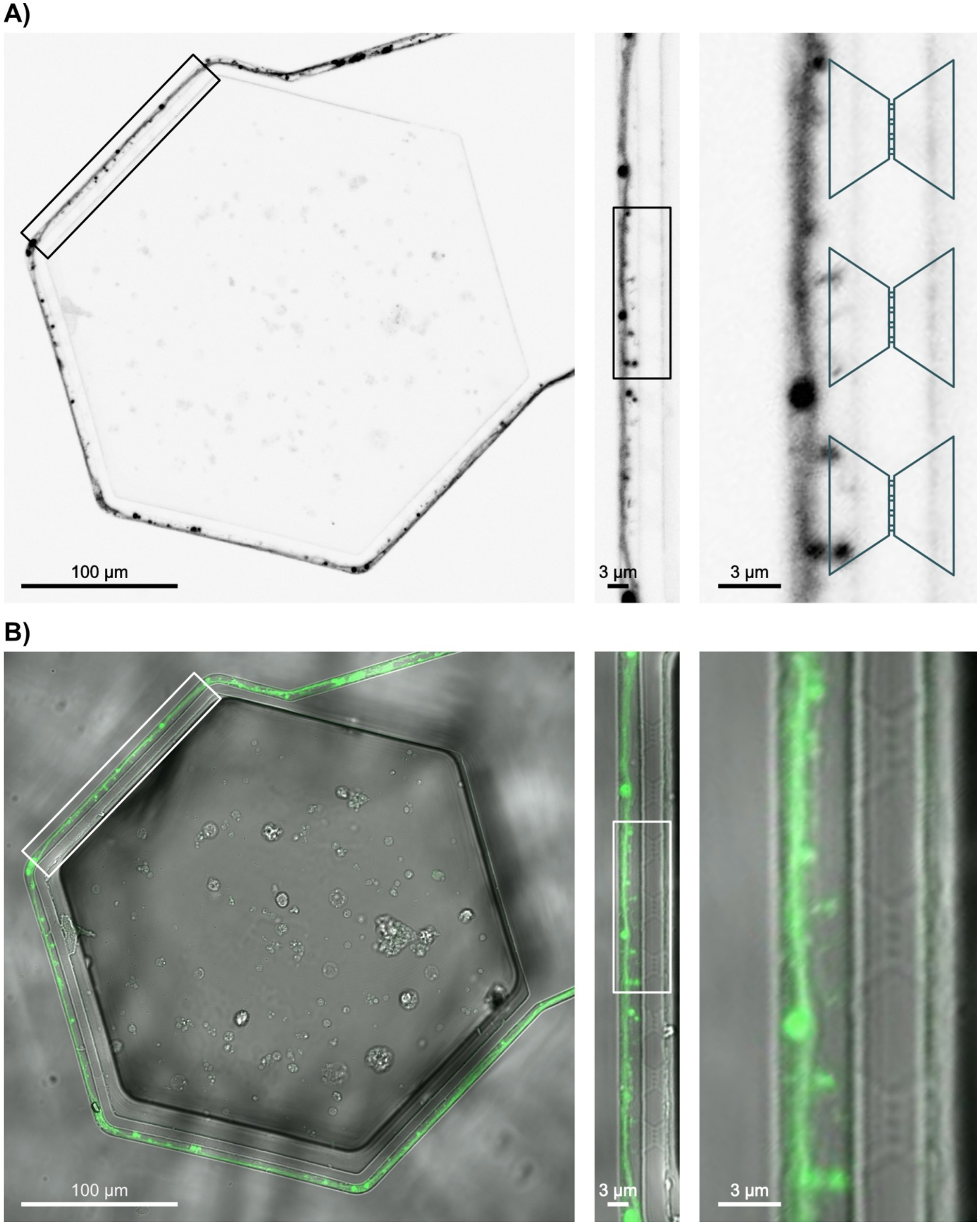


**Figure S4**: **Zoom-in of presynaptic axon restriction. A)** Same image as in **Fig. 4B** focused on the spine crossing region. **B)** Combined brightfield and fluorescence showing that the nanochannels are present and intact.

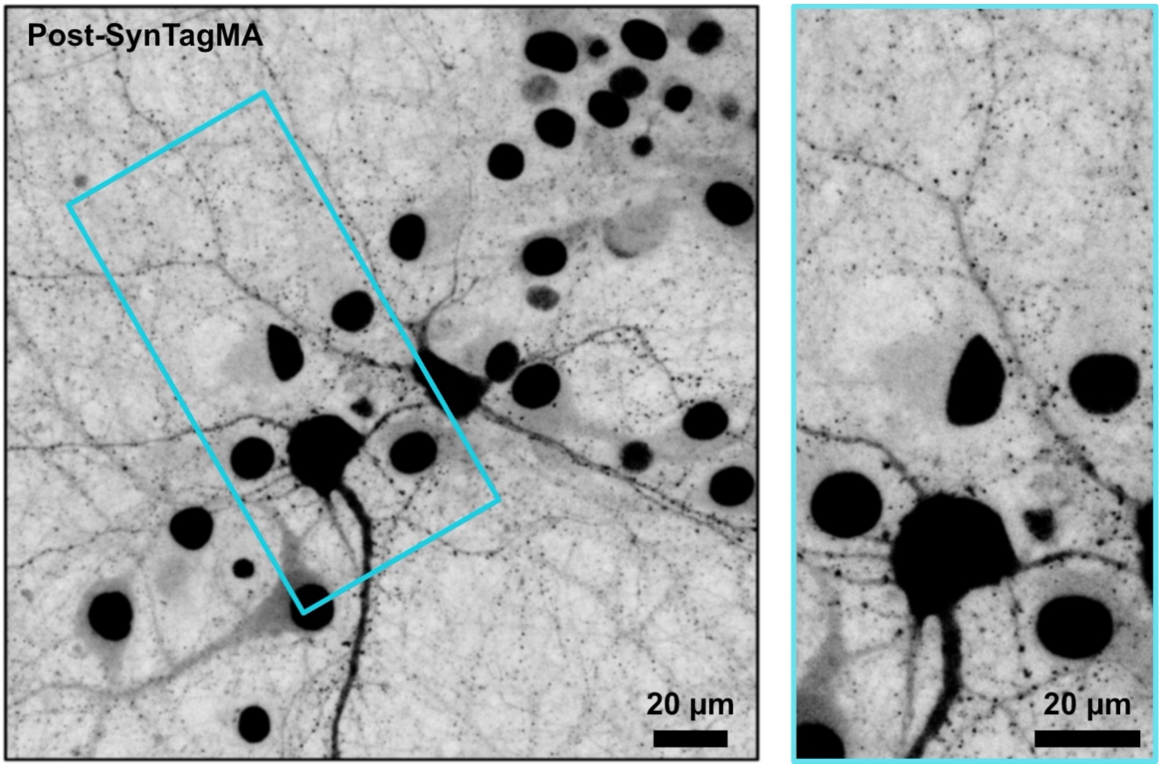


**Figure S5:** PSD95 Expression in a random culture. The image above shows that even in a small area there can be hundreds to thousands of putative synapses in a random culture. Brightness was adjusted to make puncta visible.

#### Detailed Methods for Master Mold Fabrication

Fresh Si wafer preparation:
180s Oxygen plasma, 300W
Bake on hot plate 200C for 300s

PECVD $SiO_2$:
recipe -> van Swaay $SiO_2$ 360 nm
25% $SiH_4$ ($N_2$) 500 ccm
$N_2O$ 710 ccm
LF 30 pulse 10
RF 30 pulse 10
Table heater 300C
Duration 6 min 36 sec
Measure layer using ellipsometer

EBL Layer 0; wafer prep:
300s Oxygen plasma, 600W
Bake in hot plate 200C for 300s

EBL Layer 0; spincoating:
Resist -> AR-N 7520.18, aim 360 nm
step 1 -> 500rpm, acceleration 100 rpm/s, duration 5s
step 2 -> 6000rpm, acceleration 2000 rpm/s, duration 60s
soft bake on hot plate 85C for 60s

EBL Layer 0; lithography:
Exposure performed by technician (IBM BRNC)
Develop in AR300-47 diluted 4:1 in $H_2O$, ~60s
Rinse DI $H_2O$, dry $N_2$ gun

RIE Layer 0; step 1:
recipe -> 000_SiO2 CHF3 200W
Tool -> Oxford NGP 80
$CHF_3$ 49 ccm
$O_2$ 1 sccm
RF fwd 200W, Amu 1, magnitude & phase 100%
Temperature 20C
APC 15 nTorr
LP strike 30 mTorr, 100V, ramp 5
Duration 24 min

RIE Layer 0; spincoating:
Resist -> AZ6612, 4000rpm for 40s

Expose only alignment mark region; cover structures with black film
Develop in AZ400K 4:1 in H2O

RIE Layer 0; step 2:
recipe -> 000_Si etch isotropic
Duration 2 min

RIE Layer 0; resist strip:
Acetone to strip AZ6612
PGMEA to strip AR-N 7520.18
Oxygen plasma 600W for 300s to clean surface
Inspect etched structure on microscope and profilometer

Lithography Layer 1; wafer prep:
600s Oxygen plasma, 300W
Bake in hot plate 200C for 300s

Lithography Layer 1; spincoating:
Resist -> SU8 2002, aim 20 microns
step 1 -> 500rpm, acceleration 100 rpm/s, duration 5s
step 2 -> 3000rpm, acceleration 300 rpm/s, duration 30s
soft bake on hot plate 35C for 300s, 65C for 60s, 95C for 900s
Remove resist from alignment marks using mrDev-600 developer and Q-tip
bake at 65C for 60s to dry out developer residue
Let coated wafer rest overnight

Lithography Layer 1; exposure:
Tool -> Süss MA6
Vacuum contact, intensity controller set to constant power
Mask substrate-> soda-lime glass
Mask technology -> Chromium; written by DWL
Filter -> Hoya LP360
Exposure dose -> 80 mJ/cm2, pulsed exposure 10 x 10% with 10s pause
Post-exposure bake on hot plate -> 60s at 65C, 900s at 95C

Lithography Layer 2; spincoating:
cover alignment marks with Scotch tape
resist -> SU8 3050, aim 75 microns
step 1 -> 500rpm, acceleration 100 rpm/s, duration 10s
step 2 -> 1700rpm, acceleration 300 rpm/s, duration 30s
remove tape covering alignment marks
soft bake on hot plate 65C for 120s, 95C for 2700s

Lithography Layer 2; exposure:

Tool -> Süss MA6
Hard contact, intensity controller set to constant power
Mask substrate-> soda-lime glass
Mask technology -> Chromium; written by DWL
Filter -> Hoya LP360
Exposure dose -> 300 mJ/cm2, pulsed exposure 10 x 10% with 10s pause
Post-exposure bake on hot plate -> 120s at 65C, 900s at 95C
Develop in mrDev 600 developer, rinse in IPA, dry standing, dry with N2 gun

SU8 Cross-linking:
Flood exposure in UV mask aligner for 300s, no filter or mask
Hard bake -> ramp to 200 C, bake for 900s, ramp to room temperature

Inspection:
Microscope -> check SU8 structure adhesion and alignment
Profilometer -> height measurements Layer 1, Layer 2

CVD Silanization:
Place patterned wafer in a vacuum desiccator
Add a glass vial with a small drop of perfluoro-octyl trichlorosilane
Add a pH measurement strip
Close the desiccator and apply vacuum
Leave overnight
Degas the chamber again, and purge with Nitrogen
